# An insight into new glycotherapeutic in glial inflammation: Understanding the role of glycosylation from acute to chronic phase of inflammation

**DOI:** 10.1101/2021.11.18.469119

**Authors:** Vaibhav Patil, Raghvendra Bohara, Carla Winter, Michelle Kilcoyne, Siobhan McMahon, Abhay Pandit

## Abstract

Glycosylation plays a critical role during inflammation and glial scar formation upon spinal cord injury (SCI) disease progression. Astrocytes and microglia are involved in this cascade to modulate the inflammation and tissue remodelling from acute to chronic phases. Therefore, understating the glycan changes in these glial cells is paramount. Herein a lectin microarray was undertaken using a cytokine-driven inflammatory MGC model, revealing considerable differential glycosylation from the acute to the chronic phase in a cytokine-combination generated inflamed MGC model. It was found that several N- and O-linked glycans associated with glia during SCI were differentially regulated. Pearson’s correlation hierarchical clustering showed that groups were separated into several clusters, illustrating the heterogenicity among the control, cytokine combination, and LPS treated groups and the day on which treatment was given. Control and LPS treatments were observed to be in dense clusters. This was further confirmed with lectin immunostaining in which GalNAc, GlcNAc, mannose, fucose and sialic acid-binding residues were detected in astrocytes and microglia. However, this modification (upregulation of sialic acid expression) was inhibited by the sialyltransferase inhibitor which indeed modulates the mitochondrial functions. The present study is the first functional investigation of glycosylation modulation in a MGC (MGC) model which elucidates the role of the glycome in neuroinflammation and identified potential therapeutic targets for future glycol-therapeutics in neuroinflammation

## Introduction

Central nervous system (CNS) trauma such as spinal cord injury (SCI) is dominated by inflammation with the predominance of cytokines such as TNF-α, IL-1β and IL-6. These cytokines further activate astrocytes and microglia to modulate their functions based on the expression of various glycans either on the surface or inside the cell. As the injury progresses, glial cells’ activation and their characteristics change from acute to chronic phases. Most *in vitro* studies deals with only the acute phase (24 hrs) of glial cells (Alizadeh et al., 2019). However, we previously reported a mixed glial *in vitro* model to study acute and chronic inflammatory phases associated with SCI (Patil et al., 2021). Another area studied is glial scar formation after SCI and its effect on the injury’s progression. The glial scar mainly comprises cells such as scar-forming reactive astrocytes, foam cells, microglia, M1 macrophages, etc. and extracellular matrix (ECM) (Bradbury and Burnside, 2019) (Silver and Miller, 2004). Among these cells, the astrocytes are mainly responsible for the glial scar formation and secretion of ECM components (Adams and Gallo, 2018). The glial scar comprises proteoglycans and glycoproteins, further impeding neuronal regeneration (Bradbury and Burnside, 2019, Yuan and He, 2013). The glial scar’s major constituents include chondroitin sulphate (CS) and keratan sulphate (KS) proteoglycans. These proteoglycans are made up of glycans which are polysaccharides with distinct carbohydrate chains (Bradbury and Carter, 2011, Hering et al., 2020). Glycans are mainly composed of N-acetylgalactosamine (GalNAc), N-acetylglucosamine (GlcNAc), fucose, mannose, sialic acid, galactose, lactose, or other monosaccharides (Varki, 2016). Among these, hypersialylation is a characteristic feature in which sialyltransferase increases (polysialyltransferases) and catalyses different sialic acid linkages such as α-(2-3, 2-6, or 2-8) on glycan chains (Yoo et al., 2015). In vertebrates, glycans usually end with a sialic acid (sialylation) on alpha-3 or 6 position linkage (Marsico et al., 2018) which supports molecular identification.

The modification of proteins by incorporating glycans that occurs during post-translation modification in the cytoplasm and endoplasmic reticulum is called glycosylation (Kleene and Schachner, 2004). In cellular processes, glycans play a significant role in defining the functionality of the protein or lipid. Post-translation modification of proteins and lipids is critical in pathological conditions, and aberrant changes in these modifications could alter their functionality (Helenius et al., 2001, Varki and Lowe, 2009). In addition, as stated earlier, in inflammatory conditions, the activated astrocytes and microglia modulate their functions based on the expression of various glycans either on the surface or inside the cell (Thomas and Pasquini, 2018, Starossom et al., 2012). It has been noted that GlcNAc and GalNAc moieties increased after SCI and were mostly found to be associated with microglia (2020, Rolls et al., 2008) and astrocytes (Kadomatsu and Sakamoto, 2014). Upon microglial stimulation, they exhibit pro- and anti-inflammatory responses, which are well regulated by the glycans (Puigdellívol et al., 2020). These glycans interact with their ligands known as lectins. Therefore, lectins are used to study glycan expression and identify specific glycan modifications either on the cell surface or intracellularly and on proteins or lipids. Lectins are the carbohydrate-binding moieties that recognize cell surface glycan structures, and they are involved in cell-to-cell communication via lectin-glycan interactions (Belický et al., 2016). This biomarker approach was used to study the progression of inflammatory phases from acute to chronic in the mixed glial culture (MGC) model.

As mentioned in Patil et al. (2021), using an *in vitro* MGC model, proinflammatory cytokines can induce acute and chronic levels of inflammation associated with SCI. The different types of glycans and their level of expression involved in the MGC model were further considered from the acute to the chronic stages of inflammation. Lectin microarray and lectin immunostaining were also carried out to identify the type of glycans associated with MGCs and their modulation under treatments. Notably, an attempt was made to identify overall glycosylation (O-linked and N-linked), providing insight into a new glycotherapeutic approach for understanding inflammation and glial scar formation. This phase of the study hypothesises that a cytokine-induced inflamed MGC *in vitro* model will show differential glycans from the acute to the chronic phase of inflammation associated with SCI. To investigate this hypothesis, this study included the following objectives: study the expression pattern of glycans in MGC under an inflammatory a cytokine combination and LPS treatments using lectin microarray technique; validate the glycan expression in mixed glia under a cytokine combination and LPS treatments using lectin staining. The effect of Sialyltransferase Inhibitor (STI) on sialylation expression was studied in a MGC model in an inflammatory environment to assess the efficacy of addressing glycosylation profile alterations. To the best of the authors’ knowledge, this is the first study to show that glycosylation inhibitors can replace physiological glycosylation in MGC model cells and that glycosylation plays a role in neuroinflammation. In the future, this could provide new molecular targets for biomaterial functionalization.

## Materials and Methods Cell culture

As previously described, primary MGCs were prepared from spinal cords (Kilcoyne et al., 2019). Spinal cords were isolated by hydraulic extrusion technique from three-day-old post-natal rats with minor modification in the spinal cord extrusion method (Kennedy et al., 2013). Later, meninges were gently peeled from spinal cords under microdissection microscope. Spinal tissues were chopped into fine (approx. 1 mm) pieces and digested using 1% trypsin-Ethylenediaminetetraacetic acid (EDTA) solution for 15-20 minutes. Trypsin’s activity was inhibited using Dulbecco’s Modified Eagle Medium (DMEM)-high glucose supplemented with 10 % fetal bovine serum (FBS) and 1% Penicillin/Streptomycin (P/S). Further, MGCs were cultured as previously described (Patil et al., 2021). The MGCs were grown *in vitro* for three weeks before treatments were applied. Early passage number was maintained throughout all experiments.

### Western blotting

Cells of density 5 × 10^5^ cells/mL seeded into 6-well plates and after respective treatments, cells were lysed in a radioimmunoprecipitation assay (RIPA) buffer (50 mM Tris-HCl, pH 8.0, 150 mM NaCl, 0.02% sodium azide, 0.1% sodium dodecyl sulphate (SDS), 1% Nonidet P-40, 0.5% sodium deoxycholate) (Sigma-Aldrich®, R0278) with protease inhibitor cocktail (1:100) (cOmplete™, EDTA-free, Roche, Inc., 11873580001), phenylmethylsulfonylfluoride (1:50) (Sigma-Aldrich®, 93482) and phosphatase inhibitor cocktail (1:10) (PhosSTOP™, Roche, Inc., 04906845001). The protein concentration in the total cell lysate was determined using a BCA protein assay kit (Pierce™ Bicinchoninic acid (BCA) Protein Assay Kit, Thermo Fischer Scientific™, 23225). An equal amount of protein from each sample was separated by 10-12% sodium dodecyl sulfate (SDS) polyacrylamide gel electrophoresis and transferred to Hybond^®^ ECL™ nitrocellulose membrane. The membrane was blocked with either 5% milk or 5% bovine serum albumin (BSA) depending upon the suitability of primary antibodies to avoid non-specific binding of antibodies. This was followed by primary antibody incubation overnight at 4°C with 1:2000 rabbit anti-P-NFκB-p65 (Ser536) (93H1) (Cell signalling, 3033s), mouse anti-NFκB-p65(F-6) (Santacruz biotech, sc-8008) or one hour (hr) at room temperature (RT) with 1:15,000 anti-β-actin (Sigma, A5441) on a rocking platform. All the washing steps were carried out in Tween-20 in TBS (0.1%). Next, horseradish peroxidase-conjugated secondary goat anti-rabbit or goat anti-mouse antibodies (prepared in 5% milk or 5% BSA at 1:10,000) were applied followed by enhanced chemiluminescence detection. Signals from protein bands were captured on X-ray films (CL-XPosure™ Film, Thermo Scientific™, 34090) which were further analyzed using Image Studio™ Lite software and signals recorded as pixel density.

### Cell protein extraction and glycoprotein sample labelling

As depicted in **Figure 1**, MGCs were seeded at a density of 1× 10^5^ cells/well on PLL-coated 24-well plates with media (DMEM + 1% P/S + 10% FBS). After growing in an incubator for two days, the cytokine treatment was given for 21 days as described in (Patil et al., 2021). Cells were treated with a combination of 10ng/mL of three cytokines (TNF-α, IL-1β, and IL-6) (R & D Systems; Recombinant rat TNF-α protein, 510-RT-010; Recombinant rat IL-1β/ IL-1F2 protein, 501-RL-010; Recombinant rat IL-6 protein, 506-RL-010) at day 0 and every subsequent two days (i.e., day 0, 2, 4, 6, 8, 10, 12, 14, 16, 18 and 20) until day 21. At each of the treatment time-points (except day 0), half of the media was changed and filled with fresh cytokine treatment media. An LPS treatment group underwent the same procedure, with the cytokines substituted by a 100ng/mL dose of LPS (Sigma Aldrich). The ctrl (control) group was carried out by performing the same half-media change procedure with low-serum media containing no added inflammatory molecules.

**Figure 1.**
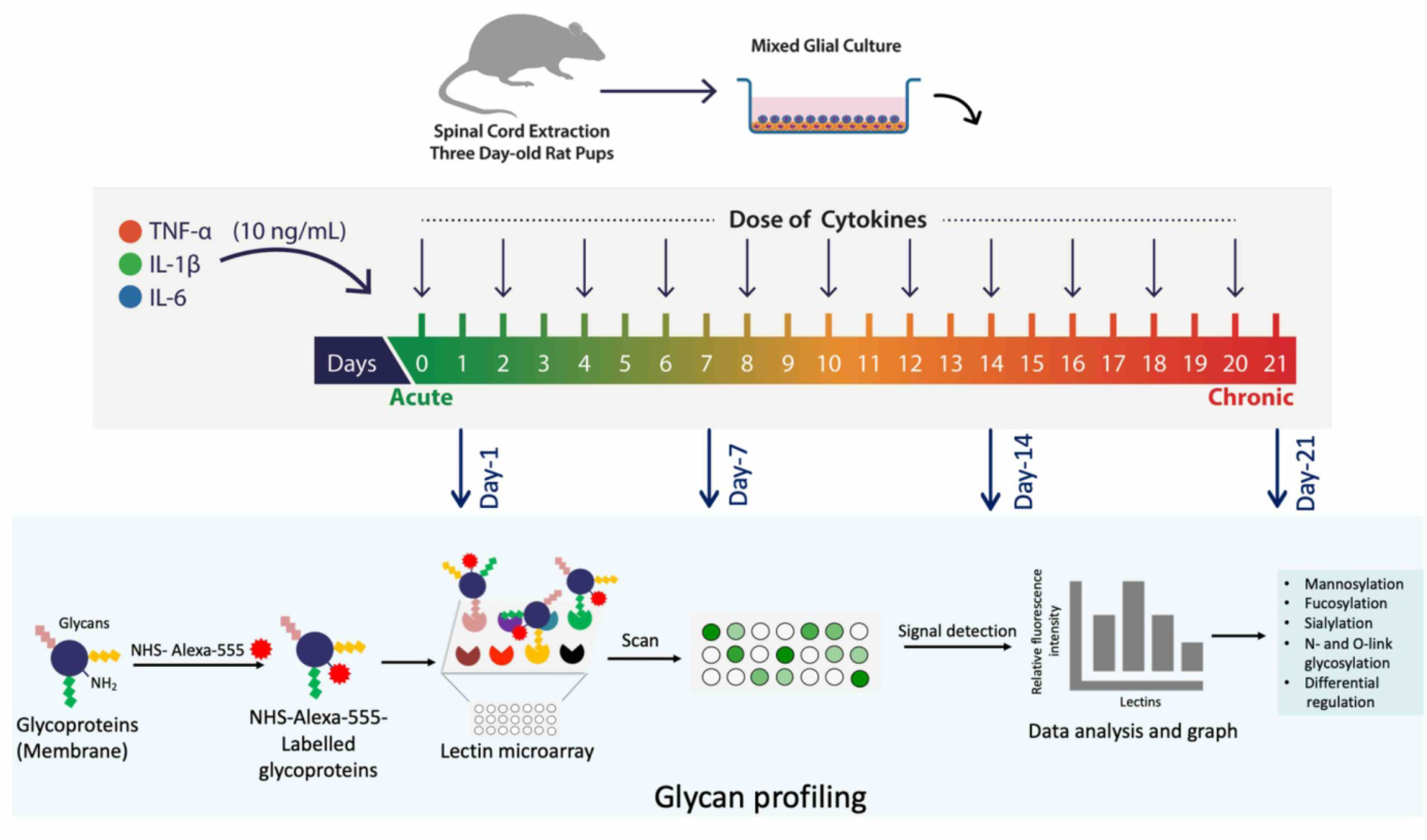
Workflow describing steps involved in the lectin microarray. MGCs were prepared from spinal cords isolated by the hydraulic extrusion technique from three-day-old post-natal rats. MGCs were treated with TNF-α, IL-1β and IL-6 (10 ng/mL per cytokine) in combination/s for one day (acute) to 21 days (chronic). Lipopolysaccharide (LPS) (10 ng/mL) was used as positive control. Treatments were given every alternate day (n=3). Membrane proteins were extracted using Mem-PER™ Plus, a membrane protein extraction kit. Lectin microarray profiling was performed using a panel of 48 lectins to study any alteration in mammalian glycosylation. MGCs were then assessed for glycosylation localisation and association with particular cell types using FITC-conjugated lectins and glial markers (GFAP and CD11b) respectively by ICC.

Samples of each group were extracted from the culture on days 1, 7, 14, and 21. A membrane Protein Extraction kit (Mem-PER ™ Plus, 89842) from Thermo Scientific was used to extract samples from cells (adherent mammalian cells). In short, after media removal cells were scraped and washed with cell wash solution. Harvested cells were permeabilised using permeabilization buffer, centrifuged, and the supernatant was collected (as it contains cytosolic proteins) and stored at -80oC. The pellet was resuspended with Solubilization Buffer, incubated, centrifuged, and the supernatant was collected as it contains membrane proteins and stored at -80°C. The membrane protein fraction would be further used for lectin micro-array.

10 μL of phosphate buffer 10x (500mM, pH=8.3) was added to the glycoprotein solutions in PBS (100μL). Afterwards, the solution was treated with Alexa-555-N-Hydroxysuccinimide (NHS) dye (1 μg/μL in Dimethyl Sulfoxide (DMSO) for 1hr at room temperature (RT) in such a way that 0.15μL of dye was employed to label 1μg of glycoprotein. The addition of Tris buffer quenched the excess of dye. The fluorescently labelled glycoprotein solutions were not purified and were directly used in the lectin array analysis.

### Lectin microarray construction, incubation, and data extraction

Lectin solutions 0.4 – 0.5 mg/mL and antibody solutions 0.1 mg/mL, were prepared in print buffer (1.0 mM D-glucose in PBS containing 0.01% of cy3-conjugated BSA). Aliquots of 20 µL volume of each freshly prepared lectin solution were loaded in a 384 well plate. Volumes (0.67 nL) of these dilutions were spotted onto an N-Hydroxysuccinimide (NHS) functionalized glass slide in 6 replicates. The humidity in the print chamber was maintained at around 50% before and during printing. The slides containing the lectin arrays **(Supplementary table 1)** were incubated after printing in a 75% humidity chamber at 18°C overnight and stored at 20°C without quenching if not used immediately. The remaining NHS groups were quenched by placing the slides in a 30 mM ethanolamine solution in borate buffer for one hr and blocked with a 0.3 mg/mL bovine serum albumin (BSA), 0.3 mM Ca^2+^ solution in phosphate-buffered saline (PBS) buffer containing 0.5% Tween-20 for one hr.

Printed slides, stored at -20°C, were quenched (NHS-activated hydrogel coated glass surface) by immersion in a 50 mM ethanolamine solution in borate buffer (50 mM, pH 8.5) for 30 mins at RT. The quenched surface was then passivated by incubation in PBS buffer containing 0.5% Tween-20, BSA 0.4 mg/mL, CaCl2 0.3 mM, for 30 minutes at RT. The glycoprotein samples were added to the corresponding wells, and the incubation was carried out for 1.5 hr at RT. Then the glycoprotein solutions were removed, and the slide washed with PBS for five mins and dried by centrifugation and scanned. The images obtained from the G265BA microarray scanner (Agilent Technologies) were analysed with Pro ScanArray Express™ software (PerkinElmer). Samples were incubated in the lectin array at a concentration of 3 µg/mL. The dilution of the samples was performed by the addition of lectin incubation buffer to the labelled glycoprotein solutions after labelling. The labelled-BSA and control dye sample was incubated under the same concentration conditions as negative controls for lectin binding. Additionally, ‘mean pixel density’ from each lectin was calculated, and the data plotted as a heatmap. Hierarchical clustering of the data was performed using Morpheus, https://software.broadinstitute.org/morpheus.

### Lectin cytochemistry

High-grade bovine serum albumin (BSA; Sigma, A7638) was treated with periodic acid (Sigma, P0430) before its use for blocking according to an established protocol (Glass et al., 1981). Briefly, 5g of BSA was dissolved in 100mL of freshly made periodic acid solution (10mM periodic acid in 0.1M sodium acetate, pH 4.5) and incubated at room temperature for six hr. The BSA was dialyzed against the water with four water changes over two days at 4ºC and finally lyophilized and stored at 4ºC. The cytochemistry procedure was followed as discussed (Rebelo et al., 2021) with few modifications for staining on cells. First, the cell media was removed, and the cells were washed twice with 1X Tris-buffered saline (TBS) (10mM Trizma hydrochloride, 0.1M NaCl, 1 mM CaCl2, 1 mM MgCl2, pH 7.2). Cells were then fixed in 4% PFA for 15 minutes, washed four times with TBS, and then blocked with 2% periodic acid-treated BSA in TBS for 30 minutes at room temperature. The cells were incubated with the appropriate concentration of fluorescein isothiocyanate (FITC)-labelled lectins (EY Laboratories Inc., San Mateo CA) in the dark for one hr at RT at the following concentrations (**Supplementary table 2**): Phaseolus vulgaris erythroagglutinin (PHA-E) and Ricinus communis (RCA)-I at 10μg/mL, Sambucus nigra (SNA)-I, Peanut agglutinin (PNA), Wheat germ agglutinin (WGA), Maackia amurensis agglutinin (MAA), Galanthus nivalis agglutinin (GNA), Datura stramonium (DSL) and Ulex europaea agglutinin (UEA)-I at 20μg/mL, and Wisteria floribunda agglutinin (WFA) at 30μg/mL. Negative controls were carried out in parallel with a TBS solution containing no lectins. Haptenic sugar inhibition controls were carried out in parallel by co-incubation of 100mM appropriate sugar in TBS with the following lectins as follows (**Supplementary Table 2**): SNA-I, WFA, and PNA were prepared in lactose, WGA and GNA in mannose, MAA and RCA-I in galactose, PHA-E in bovine IgG, DSL in N-Acetyl-D-glucosamine (GlcNAc) and UEA-I in fucose (all carbohydrates, Sigma Aldrich). After lectin staining, a traditional immunocytochemistry procedure was carried out to label astrocytes (GFAP) and microglia (CD11b).

### Immunocytochemistry

After lectin cytochemistry, cells were permeabilized by incubating in 0.2% Triton X-100 in PBS and washed with PBS. Unspecific bindings were blocked with 1% BSA+ 10% NGS in PBS solution (blocking buffer) for one hr at RT. Primary antibodies, anti-GFAP (1:500, Dako, Z033429), anti-CD11b (1:200 Sigma, CBL1512) were prepared in blocking buffer and incubated overnight at 4°C. Incubated with secondary antibody Alexa Fluor® 488 (1:500, Thermo Scientific™, A-10667) and Alexa Fluor® 546 (1:500, Thermo Scientific™, A11035) for one hr at RT. Coverslips were mounted on slides by placing a small drop of mounting medium containing DAPI (Fluoromount-G™, Thermo Scientific™, 00-4959) on the slide and placing the coverslip cell-surface down.

### Microscopy and image analysis

Fluorescent cytochemistry images were captured on an Olympus VS120 Virtual Slide Microscope with Olympus VS fluorescence software (VS-ASW-FL). Every image was taken at the same exposure fluorescence intensity to ensure consistency. Cell morphology and lectin-localization were qualitatively assessed for each sample. Additionally, lectin staining fluorescence intensity (FI) was quantitatively analyzed using the ImageJ (Fiji) software. The number of Hoechst-positive nuclei was quantified. Fluorescence intensity per cell (FIPC) was calculated by dividing the FI by the number of Hoechst-stained nuclei per image. FIPC was measured for each sample (n=3), for each lectin staining (SNA-I, WFA, PNA, WGA, MAA, PHA-E, UEA-I, GNA, DSL and RCA-I), for each group (control, cytokine-treated and LPS treated) at 24 hr.

### Seahorse Cell Mito Stress assay

The assay was performed as explained in (Patil et al., 2021). 40,000 cells/well were seeded into the XFp Seahorse miniplates. After growing in an incubator for two days *in vitro* (DIV), the media was replaced with (DMEM + 1% P/S) and cells were treated with a combination of pro-inflammatory cytokines (TNF-α, IL-1β, and IL-6) of dose 10 ng/mL (per cytokine) and STI (Sialyltransferase Inhibitor, 3Fax-Peracetyl Neu5Ac) (300 µM), 566224, Sigma-Aldrich at day 0. On the day of assay, the media was replaced with glucose, pyruvate, L-glutamine and phenol red-free Seahorse XF base medium. Assay medium was prepared with the addition of glucose (25 mM), sodium pyruvate (1.0 mM) and L-glutamine (2.0 mM). The pH was adjusted using 1N NaOH to 7.4 then media filtered through a 0.2 mm filter. Before the assay, the sensor cartridge was hydrated overnight, and seahorse instrument was turned on at least five hrs before the assay. Oxygen consumption rate (OCR) and extracellular acidification rate (ECAR) was measured on XFp Seahorse analyser with the sequential addition of 1.Oligomycin (1 uM), 2.FCCP (2 uM), 3. Rotenone/ antimycin (0.5 uM) for 110 minutes with five readings per cycle.

After assay was finished, cells were lysed in a radioimmunoprecipitation assay (RIPA) buffer (50 mM Tris-HCl, pH 8.0, 150 mM NaCl, 0.02% sodium azide, 0.1% sodium dodecyl sulphate (SDS), 1% Nonidet P-40, 0.5% sodium deoxycholate) (Sigma-Aldrich®, R0278) with protease inhibitor cocktail (1:100) (cOmplete™, EDTA-free, Roche, Inc., 11873580001), phenylmethylsulfonylfluoride (PMSF) (1:50) (Sigma-Aldrich^®^, 93482) and phosphatase inhibitor cocktail (1:10) (PhosSTOP™, Roche, Inc., 04906845001). The protein concentration was determined using a BCA protein assay kit (Pierce™ BCA Protein Assay Kit, Thermo Scientific™, 23225). OCR and ECAR data were normalised on Wave software using protein concentration per well. The extracellular acidification and rate of ATP production by glycolysis and oxidation were calculated using ECAR and OCR values (Mookerjee et al., 2015).

### Statistical analysis

All statistical analyses were performed using GraphPad Prism 8.00 software. Most data were analysed by one-way analysis of variance (ANOVA) followed by Tukey multiple comparison test for comparing more than three samples, and two-tailed unpaired *t*-tests for comparing two samples with 95% confidence. *p*<0.05 was considered to be statistically significant.

## Results

The study aimed to decipher glycan modulation in glial cells involved during neuroinflammation associated with SCI. However, similar outputs could be liked with other neuroinflammatory phenomena. The study was divided into various levels to understand glycosylation in astrocytes and microglia in MGCs. Also, we focussed our approach to show the effect of the sialyltransferase inhibitor on sialic acid expression in MGC and mitochondrial respiration.

### Study of lectin microarray upon cytokine combination treatment from the acute to the chronic phase of inflammation

During glial activation, glycans are significantly expressed and modulate an immune response. Therefore, it was essential to analyse the type of glycans involved and their correlation with cell types in MGC from the acute to the chronic phase. To understand this, the membrane proteins were extracted after the treatments at days 1, 7, 14 and 21 from MGC. Each sample with glycoproteins was tagged with Alexa-555, and 3 ug/mL of it was incubated with a lectin microarray panel of 48 lectins. The signal intensity was detected to quantify the amount of glycoprotein binding with the respective lectin (**Figure 1**).

Pearson’s correlation hierarchical clustering metric analysis was performed to identify trends in the treatment groups. As depicted in **Figure 2A**, the groups were separated into several clusters, illustrating the heterogenicity among the control, cytokine combination, and LPS treated groups and the day on which treatment was given. There was a clear distinction between cytokine combination and LPS treated samples compared to the control group. The treatment groups in early time points (day one and seven) were separated into different cluster than later time points (day 14 and 21). As mentioned in the heatmap, several lectins such as NPL, GNA, LEL, STL, DSL, RCA-I, SNA, PSA, AAL, JAC and MOA showed strong binding. Therefore, further analysis was carried out to show their precise expressions.

**Figure 2.**
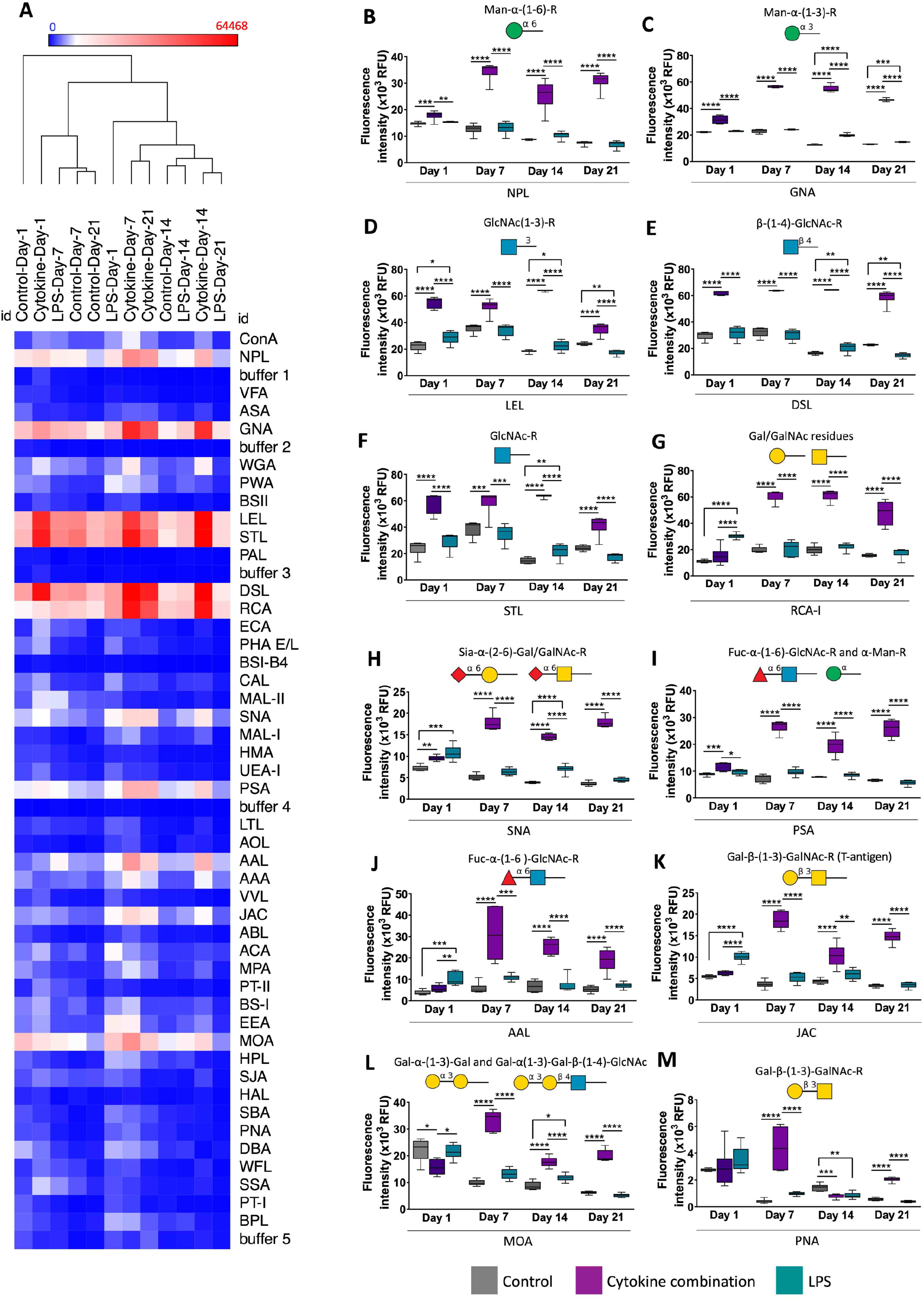
Identification of glycan modulations during cytokine combination and LPS treatments. (A) Hierarchical clustering analysis of 48 lectins. Colours define activation as highly expressed (red) and no expression (blue). Only one cytokine combination (i.e., TNF-α, IL-1β and IL-6 combination) was used along with LPS as a positive control. Treatment was given from day-1 up to day-21, and at four time points, day-1, day-7, day-14 and day-21 lectin microarray was performed on membrane proteins. Lectin microarray interaction pattern of control, at day 1, day 7, day 14 and day 21. Alexa-555-NHS dye tagged membrane glycoproteins were printed on an NHS-activated hydrogel-coated glass surface. Samples at a concentration of 3 µg/mL were incubated in the lectin array. Images were obtained by G265BA microarray scanner (Agilent Technologies) and were analyzed with Pro Scan Array Express™ software (PerkinElmer). Labelled-BSA and control dye samples were incubated at the same concentration conditions as those of negative lectin binding controls. The cytokine combination group showed an increased signal for the lectins with a higher response. In (B) Narcissus Pseudonarcissus (NPL), (C) Galanthus nivalis agglutinin (GNA), (D) Lycopersicon esculentum (LEL), (E) Datura stramonium (DSL), (F) Solanum tuberosum (STL), (G) Ricinus communis agglutinin-I (RCA-I), (H) Sambucus nigra (SNA), (I) Pisum sativum (PSA), (J) Aleuria aurantia lectin (AAL), (K) Jacalin (JAC), (L) Marasmium oreades agglutinin (MOA) and (M) Peanut agglutinin (PNA) (days 1, 7, 14 and 21), indicating higher general glycosylation upon cytokine combination treatment. Data are represented as mean ± SD, n= three independent experiments pulled samples run for six technical replicates, \**p*<0.05, \*\**p*<0.01, \*\*\**p*<0.001 and \*\*\*\**p*<0.0001. Man: Mannose, GlcNAc: N-acetylgalactosamine, Gal: Galactose, LacNAc: N-acetyl lactosamine, Sial: Sialic acid, Fuc: Fucose, α-Gal: α-Galactose.

The dye-tagged glycoproteins showed a strong interaction with mannose binding lectins such as NPL and GNA, revealing mannoside N-glycans and/or O-mannosylation (**Figure 2B-C**). MGCs also present a firm binding with N-acetyl glucosamine (GlcNAc) binding lectins such as LEL and STL, which reported the presence of multi-antennary N-glycans (**Figure 2D and 2F**). Further, the DSL lectin interaction showed the potential poly-N-acetyl lactosamine (LacNAc) epitope on the glial membrane (**Figure 2E**). The membrane glycoproteins showed binding to galactose lectins such as RCA-I, which interacts with terminal beta-galactose residues (**Figure 2G)**. There was a low interaction with ECA, which can be interpreted as a low amount of terminal N-acetyl galactosamine (GalNAc) (**Figure 2A**).

Regarding sialylation, the samples interacted with SNA lectin, reporting 2,6-sialylation, whereas the signal for MAL-I, which interacts with 2,3-sialic acid, was considerably lower (1.34-fold, *p*<0.01) at day one. However, on days 7, 14 and 21 the interaction was higher in the cytokine-combination treated group (3.45-4.8-fold, *p*<0.0001) (**Figure 2A and 2H**). Interaction with fucose-binding lectins was also crucial with PSA, showing fucosylated biantennary N-glycans and/or high mannose glycans (**Figure 2I**), AAL and AAA, which showed the presence of fucose, mainly 1,6-fucosylation (**Figure 2A and 2J**). There was also significant binding to T-antigen and α-Gal lectins, indicating the potential presence of O-glycans. The lectins JAC, ACA and MPA show the presence of T-antigen, Gal-β-(1-3)-GalNAcα O-glycan epitopes (**Figure 2A and 2K**). The lectins MOA and EEA recognize O-glycans with the Gal-α-1,3-Gal epitope. Regarding GalNAc lectins, the sample showed a signal for them at each time point, however the binding affinity was significantly reduced compared with others (3-10-fold) (**Figure 2A and 2L**).

When treatments were grouped according to days (1, 7, 14 and 21), it was observed that the signal had increased for the lectins with a higher response in the group of cytokine combination. This phenomenon was observed, for example, in the case of NPL, GNA, LEL, STL MOA, PSA, RCA-I and DSL (days 1, 7, 14 and 21) and SNA, AAL, AAA, PNA and JAC (days 7, 14, 21), indicating higher (*p*<0.0001) general glycosylation in the group mentioned above. Interestingly, the cytokine combination group showed more significant glycosylation profile (*p*<0.0001) changes than control and LPS treated groups (**Figure 2A**).

### Localisation of glycans on astrocytes and microglia

After the lectin microarray, lectin immunostaining was performed to identify the glycans’ localisation on glia and quantify the total lectins’ total intensity. GNA lectin staining showed expression of α-1,3 mannose on both astrocytes and microglia, and the LPS treated group showed higher expression at day one (7.34-fold) (**Figure 4A-4B**). PHA-E staining, which identifies GlcNAc residues, did not display any difference between control and cytokine-combination treated samples; however, LPS treatment reduced expression level (*p*<0.05) (**Figure 3I-3J**).

**Figure 3.**
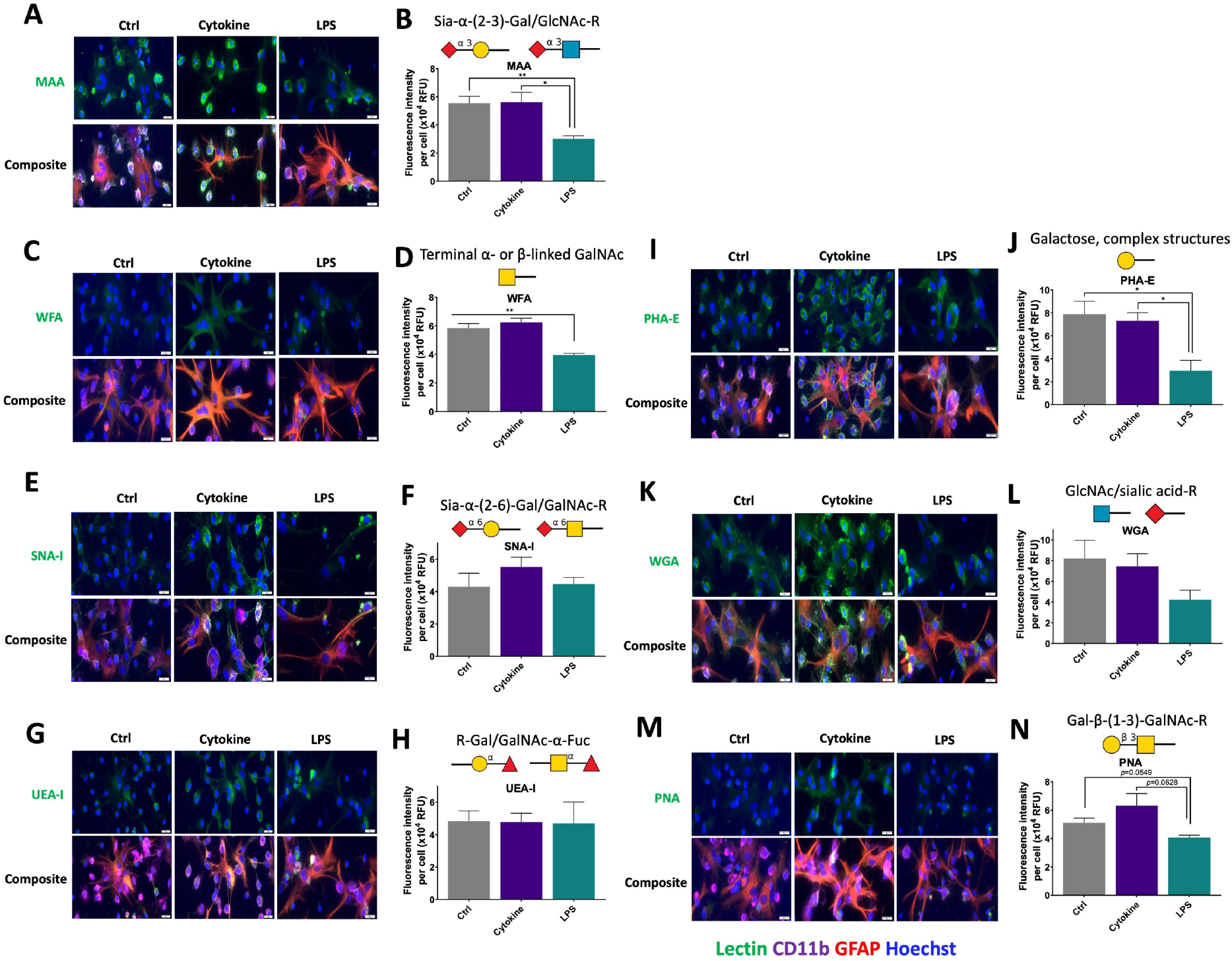
Effect of cytokine combination and LPS on the expression of lectins. Fluorescence intensity per cell was quantified and averaged for each lectin of the three treatment groups at 24 hrs. (A, B) MAA, (C, D) WFA, (I, J) PHA-E and (M, N) PNA showed significant differences between groups. (E, F) SNA-I, (G, H) UEA-I and (K, L) WGA showed no significant differences between the three treatment groups. Data are represented as mean ± SEM, n=3 three independent experiments, one-way ANOVA followed by multiple comparisons Tukey post hoc test. *p<0.05. Scale bar: 20 µm.

**Figure 4.**
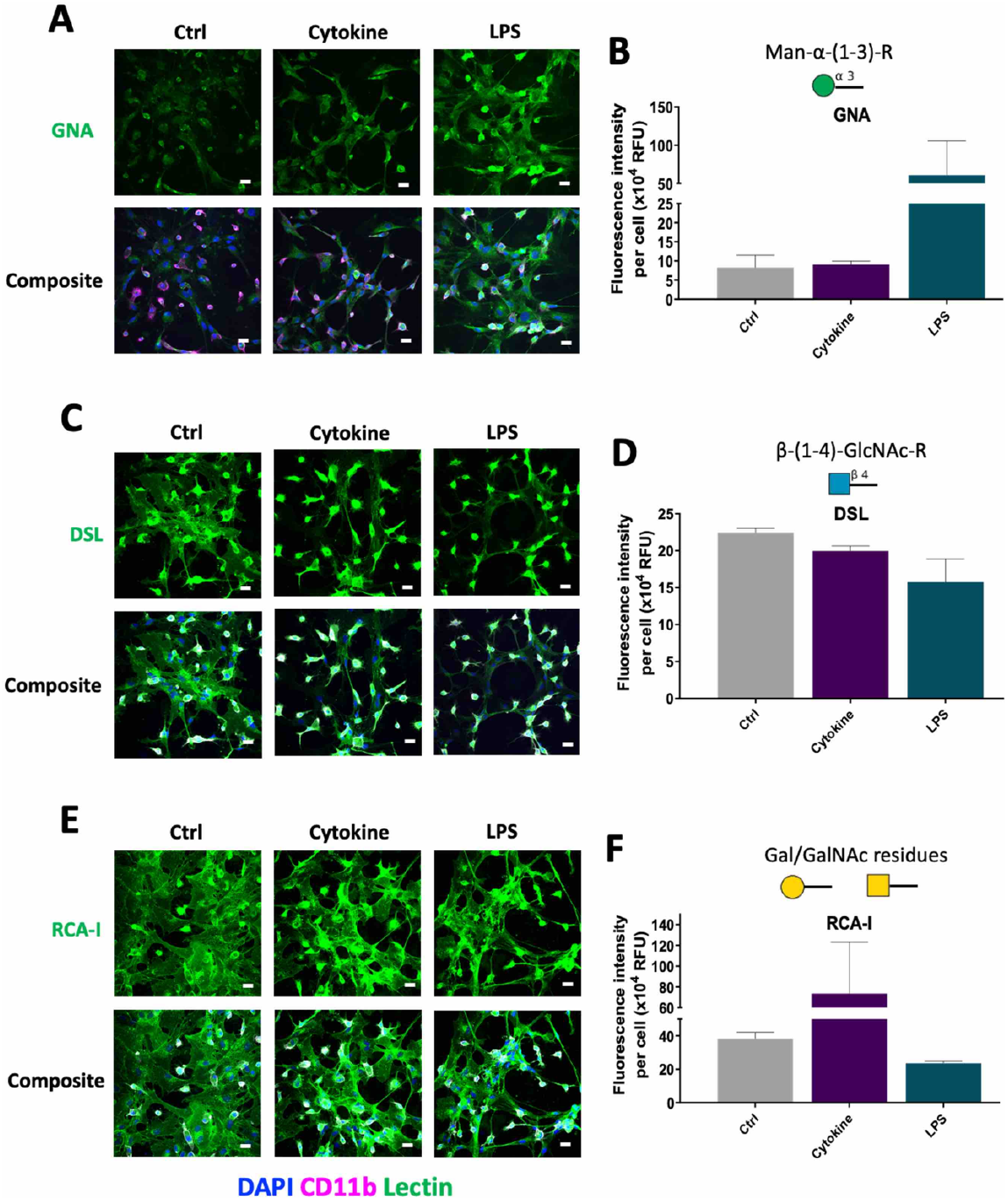
Effect of cytokine combination and LPS on the expression of glycans. Fluorescence intensity per cell was quantified and averaged for each lectin of the three treatment groups at 24 hrs. (A-B) GNA, (C-D) DSL, (E-F) RCA-I showed no significant differences between the three treatment groups at 24 hrs. Data are represented as mean ± SEM, n= three independent experiments, one-way ANOVA followed by multiple comparison Tukey post hoc test. Scale bar: 20 µm.

Upon performing DSL lectin staining (which binds β-(1-3)-GalNAc residues), it was observed that β-(1-3)-GalNAc residues were associated with microglia and astrocytes in MGC, and there was no significant difference between control, cytokine combination and LPS treated groups (**Figure 4C-4D**). Upon lectin ICC, RCA-I’s binding with GalNAc on astrocytes and microglia was observed and was higher (1.92-fold) in the cytokine-combination-treated group (**Figure 4E-4F**). WGA lectin which binds with β- (1-4)-GlcNAc residue showed increased binding with astrocytes and microglia upon cytokine combination (0.91-fold) whereas this decreased after LPS treatment (WGA, 1.95-fold) (**Figure 3K-3L**). PNA, which binds with β-(1-3)-GalNAc residue, showed increased binding with astrocytes and microglia upon cytokine combination (1.24-fold), whereas this decreased after LPS treatment (*p*<0.0549) (**Figure 3M-3N**). Further, staining was also carried out for α or β-GalNAc residues associated with chondroitin sulphate using WFA lectin. It was found that WFA binding was associated with astrocytes in the control and cytokine treated groups. However, LPS treatment showed lower WFA binding (*p*<0.01) (**Figure 3C-3D**).

Upon lectin staining, SNA-I lectin showed more binding with α-2,6-sialic acid moieties of microglia than with astrocytes, and quantification showed a higher signal in the cytokine-combination treated group (1.29-fold). LPS did not show any change compared to the control group (**Figure 3E-3F**). Additionally, MAA which identifies α-2,3-sialyation, showed strong binding with microglia and upon LPS treatment the binding was reduced (*p*<0.01) (**Figure 3A-3B**). Lectin staining using UEA-I lectin showed binding for fucosylation on both astrocytes and microglia (**Figure 3G-3H**). The summary of lectin staining outputs is mentioned in **Supplementary Table 3**.

### Sialyltransferase Inhibitor, 3Fax-Peracetyl Neu5Ac (STI) treatment downregulates the sialic acid expression

One of the glycan, sialic acid, is found to be associated with modulation of inflammation in various conditions. As stated above lectins binding with sialic acid linkages were detected in the lectin microarray, which was further confirmed by staining of MAA and SNA-I. In neural tissue, sialic acid is pre-dominantly present in the form of polysialic acid associated with the neural cell-adhesion molecule (NCAM). In CNS N-glycans are either or both sialylated and/or fucosylated, which governs a particular function of N-glycans. Hypersialylation is a characteristic feature in which sialyltransferase (along with it other 36 enzymes) increases (polysialyltransferases) and catalyzes different sialic acid linkages such as α-(2-3, 2-6, or 2-8) on glycan chains (Yoo et al., 2015). Therefore, we decided to understand the effect of Sialyltransferase Inhibitor on sialic acid expression in MGC and metabolomics associated with it under normal and inflammatory conditions.

First the dose of the drug Sialyltransferase Inhibitor, 3Fax-Peracetyl Neu5Ac (STI) was optimised using three different concentrations 100µM, 200µM and 300µM. We treated it on STI was added to MGC for seven days (in which dose was given on day 0 and day four) and upon increase in the concentration, the binding affinity of MAA lectin reduced (**Supplementary figure 1**). A dose of 300µM concentration was chosen for further use. It was observed that even three days of treatment (in which dose was given on day 0) there was decrease in MAA lectin binding (**Supplementary figure 2**). When MGC were treated for seven days with cytokine combination treatment (**Figure 5A**), there was an increase in the MAA binding intensity in the cytokine combination treatment group, which was reduced upon STI treatment, which confirmed that STI can downregulate higher expression of sialic acid duting inflammation (**Figure 5B-5F**). As NFkB-p65 is major regulator in the inflammation we studied its relevance here and performed western blotting to see the effect of STI on NFkB-p65 pathway in MGC under cytokine combination treatment. It was found that there was higher expression of this pathway in all the groups treated in combination with cytokines compred to vehicle and STI only treated groups. However, between cytokine only and cytokine with STI groups there was no significant difference (**Supplementary figure 5**).

**Figure 5.**
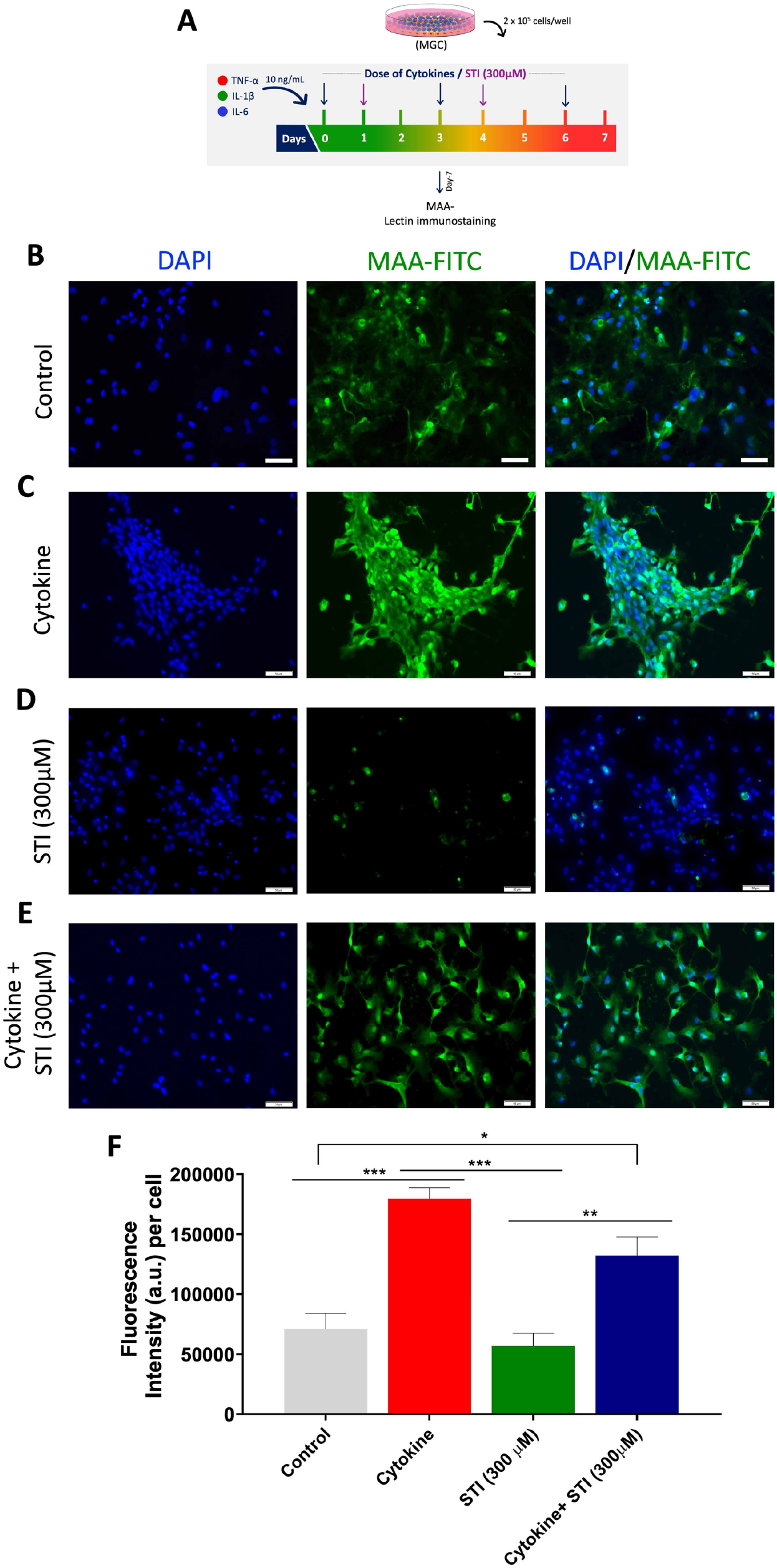
Effect of Sialyltransferase Inhibitor, 3Fax-Peracetyl Neu5Ac (STI) on MAA lectin binding on MGC under cytokine combination treatment. (A) Experimental design: 2 × 10^5^ cells/mL of MGC were seeded in 24 well plates. After 48 hrs they were given a cytokine combination treatment and this treatment was repeated on day three and day six. In control group, instead of cytokine combination treatment FBS-free DMEM media was added. Meanwhile, STI with 300 µM concentration was given on day one and subsequently on day four. At the end on day seven cells were stained with MAA lectin to see the expression of sialic acid. MAA recognizes α-(2,3)-linked sialic acid. (B) Control group, (C) Cytokine (cytokine combination), (D) STI (300 µM), (E) Cytokine + STI (300 µM) and (F) quantification of MAA binding fluorescence intensity (a. u.) per cell. Data are expressed as mean ± SEM, n= three independent experiments. One-way ANOVA followed by multiple comparison Tukey *post hoc* test. \**p*<0.05, \*\**p*<0.01, \*\*\**p*<0.001. Scale bar: 50 µm.

### Sialyltransferase Inhibitor, 3Fax-Peracetyl Neu5Ac (STI) treatment alters the mitochondrial function

Seahorse Mito Stress assay was performed on day one and day three to assess mitochondrial function in MGC when treated with STI and/or cytokine combination. Oxygen consumption rate (OCR) and extracellular acidification rate (ECAR) were measured with the sequential addition of 1. Oligomycin (1 uM), 2. FCCP (2 uM) and 3. Rotenone/ antimycin (0.5 uM). A decrease in the OCR and ECAR was observed in the cytokine-treated group at day one (**Supplementary figure 3**) but this was increased at day three (**Figure 6**), this is due to activation of the glia cells. OCR values were used to determine mitochondrial activity. To understand bioenergetics phenotypes OCR and ECAR values were used to calculate the proton production rate (PPR) and rate of ATP production by glycolysis and oxidation.

**Figure 6.**
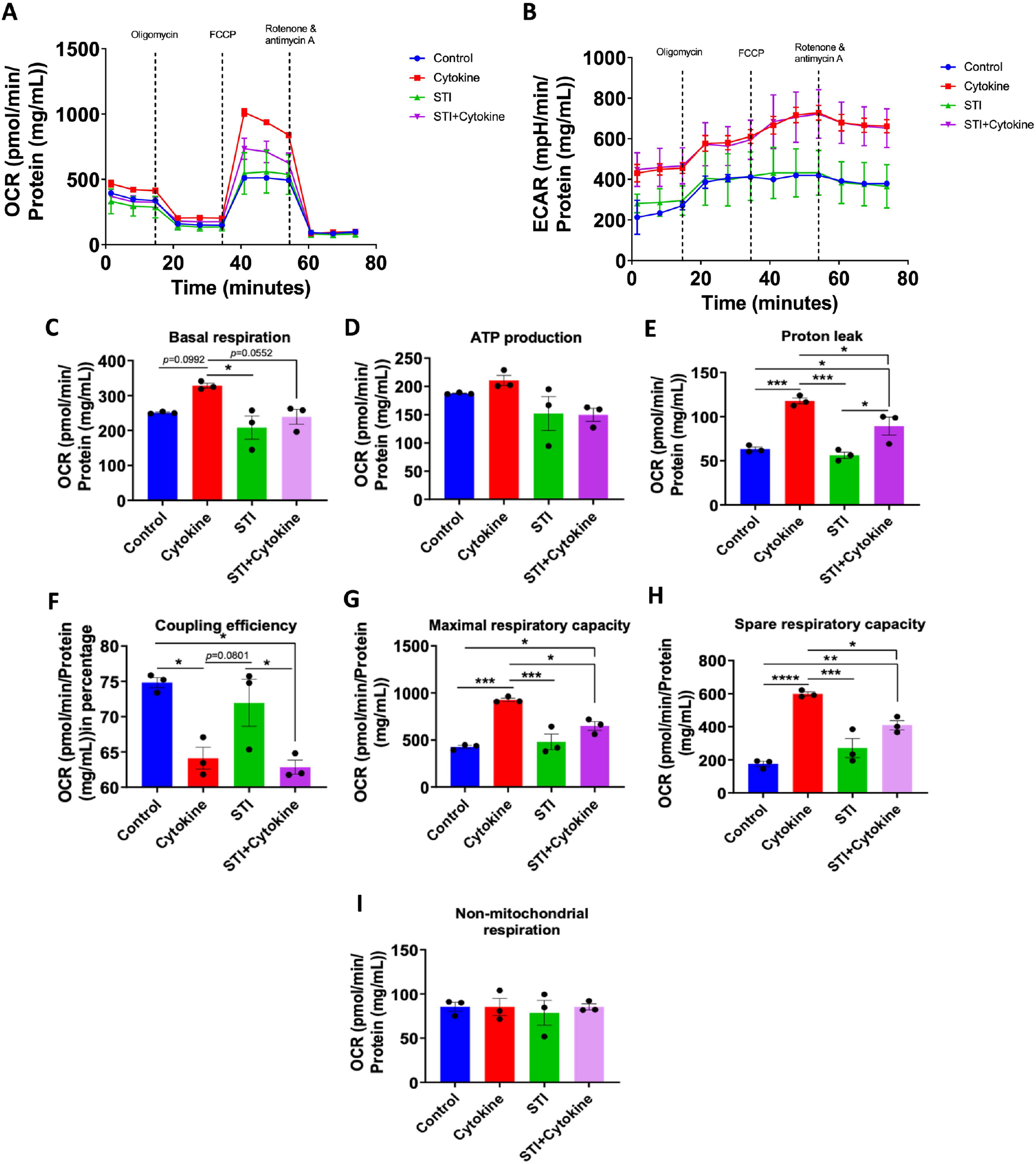
A STI (Sialyltransferase Inhibitor, 3Fax-Peracetyl Neu5Ac) treatment reverts basal respiration, ATP production, proton leak, coupling efficiency, maximal respiratory capacity and spare respiratory capacity caused by cytokine treatment by day three. After 48 hrs, MGC cells were treated with cytokine combination, with or without STI at day 0 only and on day three cells were undergone Cell Mito Stress assay. (A) OCR after the addition of three drugs (i.e. oligomycin, FCCP and Rotenone and antimycin A) sequentially. (B) Extracellular acidification rate (ECAR) after the addition of the above mentioned three drugs sequentially. (C-I) All parameters were calculated as a function of a cytokine combination and STI treatment. For this, total protein per well was calculated using a BCA protein quantification assay and data was normalised against it. Seven parameters namely, basal respiration, ATP production, proton leak, coupling efficiency, maximal respiratory capacity, spare respiratory capacity and non-mitochondrial respiration were measured and plotted as a bar graph. Data are represented as mean ± SEM, n=3. \**p*<0.05, \*\**p*<0.01, \*\*\**p*<0.001, \*\*\*\**p*<0.0001. One-way ANOVA followed by multiple comparison Tukey *post hoc* test was performed.

The first parameter examined was basal respiration and it was found that control and STI treated groups had lower basal respiration than cytokine treated group at day three (**Figure 6A**). Wheras upon STI treatment it was decreased even in combination with cytokine at day one and three (**Figure 6C and Supplementary figure 3C**). This signifies that STI was able to reduce activation of the glial cells and it had effect on mitochondial activity. This can be correlated with the acidification by oxidation parameter, as STI was able to reduce the acidification by oxidation (**Figure 7B**). Similar outputs were also observed with ATP production as basal respiration is directly linked with ATP production (**Figure 6D and Supplementary figure 3D**). Further analysis showed that the rate of ATP production by glycolysis was higher in cytokine treated groups compared to rate of ATP production by oxidation (**Figure 7C**). Later the proton leak, which is a parameter often considered a hallmark of mitochondrial damage, was calculated and found that it was higher in the cytokine group and it was significantly reduced upon STI treatment at day three (**Figure 6E**). In contrast, the coupling efficiency in these two groups was lower compared to the control and STI only treated group at day three (**Figure 6F**). The latter finding was expected as cytokine treatment imbalances the mitochondrial ability to couple number of protons and the amount of ATP produced. The FCCP dependent maximal respiration (**Figure 6G**) and spare respiratory capacity (**Figure 6H**) was higher in the cytokine treated group and it was reduced upon STI treatment which confirms that STI has an overall effect on astrocytes and microglia to lower their activation.

**Figure 7.**
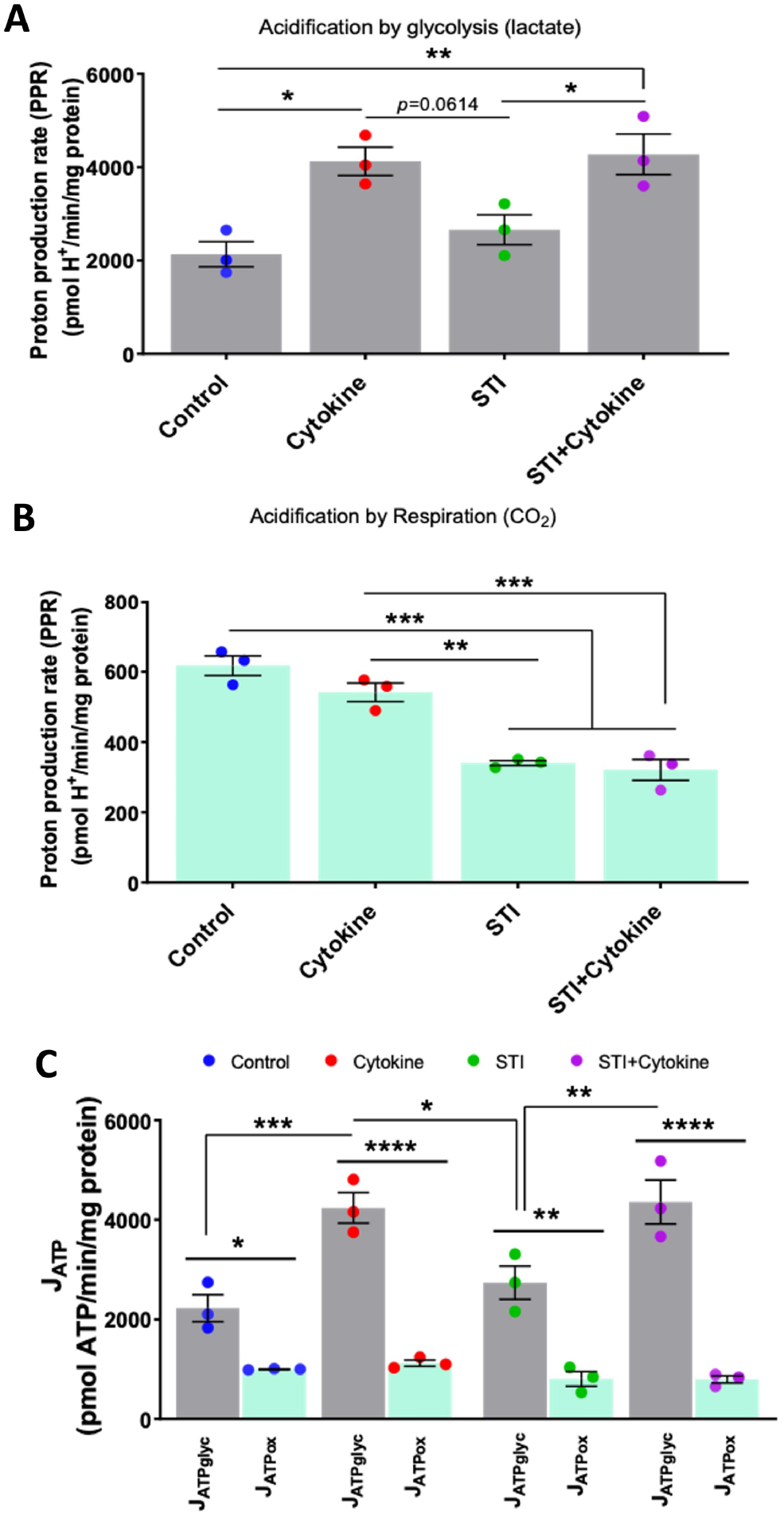
Bioenergetics phenotype of mitochondrial function (at day three). After 48 hrs, MGC cells were treated with cytokine combination, with or without STI at day 0 only and on day three cells were undergone Cell Mito Stress assay. (A) The rate of extracellular acidification caused by glycolysis due to lactate production. Both the cytokine and the STI + cytokine treatment groups showed an increase in acidification by glycolysis compared to the control and the STI group. (B) The rate of extracellular acidification due to respiration by CO_2_ production. It was higher in control and cytokine treated group compared to the STI and STI + cytokine treated groups. However, there was no difference between the STI and the STI + cytokine treatment group in acidification by respiration. (C) Data from basal ECAR and OCR has been converted to the rate of ATP production by glycolysis and oxidation using formula. The rate of ATP production by glycolysis was higher in the cytokine and the STI + cytokine treatment groups compared to the control and the STI group (J_ATPglyc_). Under four treatments the rate of ATP production by glycolysis was higher than the rate of ATP production by oxidation. Data are represented as mean ± SEM, n=3. \**p*<0.05, \*\**p*<0.01, \*\*\**p*<0.001, \*\*\*\**p*<0.0001. One-way ANOVA followed by multiple comparison Tukey *post hoc* test was performed.

## Discussion

The inflammation associated with SCI often linked with the activation of microglia and astrocytes (Popovich et al., 1997) as these cell types govern the regenerative capability of severed spinal cord axons. Along with inflammation another major characteristic feature of SCI is a glial scar formation because of astrogliosis (activation of astrocytes). These are the major events which are responsible for the death of healthy oligodendrocytes and neurons around the lesion site (Fitch and Silver, 2008). We have established an *in vitro* model of MGC prepared from the spinal cord of postnatal rats to study not only inflammation but also changes in glycans from acute to chronic phases.

### Differential glycosylation in MGC model from acute to chronic phases of inflammation

Glycans are polysaccharides present in all living organisms. They are fundamental building blocks of cellular structure, energy storage and system regulators. They form an integral part of several glycoconjugates such as glycolipid, glycoprotein, or proteoglycans (Wiederschain, 2013). Glycans are divided into several classes: glycosaminoglycans (GAGs), N-linked glycans, O-linked glycans, glycosphingolipids, and glycosylphosphatidylinositol (GPI) anchor (Marsico et al., 2018). GAGs are of several types, such as hyaluronan (hyaluronic acid, HA), chondroitin sulphate (CS), heparin sulphate (HS), keratan suphate (KS) and dermatan sulphate (DS) (Kleene and Schachner, 2004). N-linked glycans are processed or added into proteins/lipids in the endoplasmic reticulum to asparagine residues presents on proteins through an N-acetylglucosamine (GlcNAc). There are four types of N-glycans: high mannose, bi-antennary, tri-antennary and Tetra-antennary (Zhao et al., 2006, Vervecken et al., 2004). O-linked glycans are processed or added in the Golgi apparatus to serine, threonine, or tyrosine on proteins through an N-acetylgalactosamine (GalNAc). O-linked glycans can also be added to mannose (Man), fucose (Fuc) or glucose (Glc) residues (Szulc et al., 2020).

The O-glycosylation and N-glycosylation of several proteins defines their functions. For example, collagens, fibrin, fibronectin, laminin, HA, etc. are either N-glycosylated or O-glycosylated (Novak and Kaye, 2000). In CNS N-glycans are either or both sialylated and/or fucosylated, which governs a particular function of N-glycans. Hypersialylation is a characteristic feature in which sialyltransferase (along with it other 36 enzymes) increases (polysialyltransferases) and catalyzes different sialic acid linkages such as α-(2-3, 2-6, or 2-8) on glycan chains (Yoo et al., 2015). In vertebrates, glycans usually end with a sialic acid (sialylation) on α-3 or 6 position linkage (Marsico et al., 2018), which supports molecular identification. Another example of O-glycosylation and N-glycosylation of proteins is a family of receptors called integrins. These interact with the surrounding glyco-microenvironment and help in cell attachment and in cell-cell communication (Hynes, 2002).

Glycans are ubiquitous throughout the nervous system and play crucial roles during cellular differentiation, immune cell activation and homeostasis (Mendez-Huergo et al., 2014), (Marsico et al., 2018). Therefore, it was critical to understand the role of glycans during inflammatory conditions and their expression patterns. Glial activation from the acute to the chronic phases of inflammation have an impact on cellular dynamics and extracellular matrix (Kleene and Schachner, 2004). GlcNAc moieties are essential not only in inflammation but also in glial scar formation. They are essential for the biosynthesis of keratan sulphate which is a part of glial scar (Lind et al., 1993),(Funderburgh et al., 1991). N-linked glycosylation and mannosylation are reported to be involved in acute and chronic inflammation (Albrecht et al., 2014).

Glycans linkages expressed on glia were identified in the treatments which grouped according to days (1, 7, 14 and 21), it was observed that, in the group of cytokine combination, the signal had increased for the lectins with a higher response. This phenomenon was observed, for example, in the case of NPL, GNA, LEL, STL and DSL (day 1, 7, 14 and 21) and SNA, PSA, AAL, AAA, JAC and MOA (day 7, 14, 21), indicating higher general glycosylation in the mentioned group. The cytokine combination group would be the one that would differ from the control and LPS groups (Figure 1A-1E). We observed higher expression of sialylation, fucosylation which are involved in inflammation paradigm. In addition, N-linked acetyl glucosamine and galactosamine which takes part in the biosynthesis of keratan sulphate and chondroitin sulphate respectively, which are the main components of astrocytic scar formation after SCI. The higher sialyation upon cytokine treatment in the later stages of inflammation which confirms the (findings of) reports where higher sialyation and fucosylation was observed in response to the inflammation (Yasukawa et al., 2005). N-acetyl galactosamine polymers are essential for the synthesis of chondroitin sulphate (Silbert and Sugumaran, 2002), and therefore these glycans play a significant role during glial scar formation.

### Downregulation of sialic acid expression upon STI treatment

Because of their relative stability and ease of distribution, small-molecule inhibitors have recently received a lot of attention in biomedical applications. The ability to easily encapsulate or can be used for biomaterial functionalization, small molecules are gaining popularity. (Natoni et al., 2020). In the present research we used STI inhibitor and studied at in vitro level for their ability to alter the glycome. We also observed that these compounds had no negative effects on cell viability even at higher concentration.

### STI treatment reverts the mitochondrial dysfunction caused by a cytokine combination treatment

The primary function of mitochondria is to provide energy to the cell by producing adenosine tri phosphate (ATP), a cell’s energy currency (Wai and Langer, 2016). Apart from this, mitochondria play a role in cellular homeostasis, controlling cell growth and cell cycles, calcium homeostasis and apoptosis (Wai and Langer, 2016). Neuronal cells and glial cells rely on mitochondria’s optimum functioning, and any impairment could lead to neurodegenerative disease (Burté et al., 2015). In addition, during injury or disease conditions, malfunction in the mitochondrial activities can trigger a hyper response, which further damage healthy cells and slowing the recovery.

Microglia, upon activation, depend heavily on glycolysis for their metabolism and, astrocytes depend on glycolysis and oxidative phosphorylation for their energy generation and, upon activation, they switch to glycolytic pathways for adenosine tri phosphate (ATP) production (Hertz et al., 2007). Therefore, both cell types have a loosely assembled mitochondrial respiratory chain which could trigger higher ROS generation and promote a maladaptive glial response (Lopez-Fabuel et al., 2016, Vicente-Gutierrez et al., 2019, Ye et al., 2017). Due to activation, their metabolism increases and so does the glucose consumption. This increases lactate production via glycolysis and CO_2_ production via mitochondrial oxidative phosphorylation (respiration) which acidifies the medium.

Another study showed that under inflammation stimuli (Joshi et al., 2019) or several pathological conditions (Hou et al., 2019) mitochondrial changes causes impact on alleviating the disease conditions. Inflammation triggered in MGC caused mitochondrial respiration impairment as proton leak was increased and coupling efficiency was decreased. This was caused by a greater number of cells losing mitochondrial membrane potential upon cytokine combination treatment. Excessive inflammation in glia increases demands for ATP production which impairs the electron transport chain, therefore, increase in the glycolysis. Due to the unbalance between energy need and capacity of mitochondria there is leakage of protons from the inner membrane of mitochondria. This was caused by a greater number of cells losing mitochondrial membrane potential upon cytokine combination treatment. Excessive inflammation in the glia increases demands for ATP production which impairs the electron transport chain, and therefore an increase in the glycolysis. Due to the imbalance between energy need and the capacity of mitochondria there is leakage of protons from the inner membrane of mitochondria. In the end, this leads to the production of ROS (Visavadiya et al., 2016). This data was correlated with the proton leak parameter which determines whether there is mitochondrial membrane depolarization (Cheng et al., 2017). It is been documented that immune cells with quiescent and anti-inflammatory phenotype are primary based on fatty-acid metabolism, in contrast pro-inflammatory phenotypic immune cells use glycolysis as the main source of ATP production (Nonnenmacher and Hiller, 2018).

In conclusion, Glycosylation plays a critical role during SCI progression. In this study, glycan changes were evaluated in the mixed glial inflammatory model. First, a lectin microarray was performed which showed significant differential glycosylation from the acute to the chronic phase in a cytokine combination induced inflamed MGC model. It was found that several N- and O-linked glycans associated with glia during SCI were differentially regulated. This was further confirmed with lectin immunostaining in which GalNAc, GlcNAc, mannose, fucose and sialic acid binding residues were detected in astrocytes and microglia. The LPS treated group showed lower binding than those/that of the cytokine-combination treated group. This model can study the progression of inflammation and corresponding glycosylation changes on the glial cell surfaces. STI was able to reduce the expression of sialic acid in MGC and even along with cytokines. Mitochondrial function assay revealed that the impairment caused due to cytokine was significantly reversed by decreasing hypersialylation. This model can be used to study the progression of inflammation and corresponding glycosylation changes on the glial cell surfaces.

## Acknowledgments

We would like to thank Anthony Slone for editing the manuscript assistance in flow <u>cytometry</u>. The authors acknowledge the facilities and scientific and technical assistance of the Centre for Microscopy & Imaging at the National University of Ireland Galway (www.imaging.nuigalway.ie). We are grateful to the facilities and scientific and technical assistance of the Genomics and Screening and Flow Cytometry Core at the National University of Ireland Galway, a facility that is funded by the Science Foundation Ireland, National University of Ireland, Galway and the Irish Government’s Programme for Research in Third Level Institutions, Cycles 4 and 5, National Development Plan 2007-2013, and the European Regional Development Fund. This publication has emanated from research supported in part by a grant from the Science Foundation Ireland and is co-funded under the European Regional Development Fund under grant number <u>13/RC/2073_P2</u>.

## Notes

### Competing Interest Statement

The authors have declared no competing interest.

