## Supplementary material for "An insight into new glycotherapeutic in glial inflammation: Understanding the role of glycosylation from acute to chronic phase of inflammation": Supplimentary file

+Equal authorship

*Corresponding author

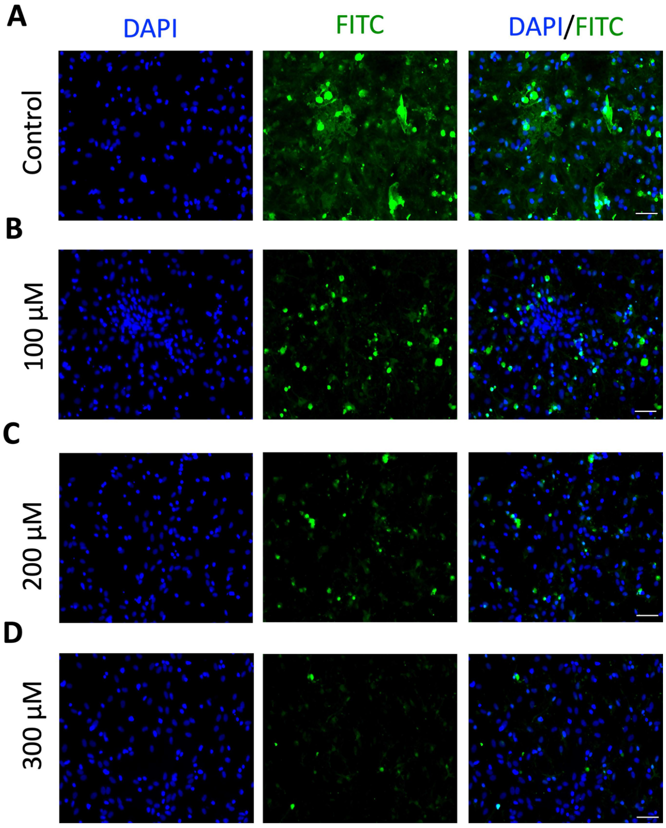


**Supplementary figure 1. Optimization of drug dose for Sialyltransferase Inhibitor, 3Fax-Peracetyl Neu5Ac (STI).** We have used three different concentrations (A) Control, (B) 100µM, (C) 200µM and (D) 300µM. Upon increase in concentration the binding affinity of MAA lectin reduces. Scale bar:50 µm.

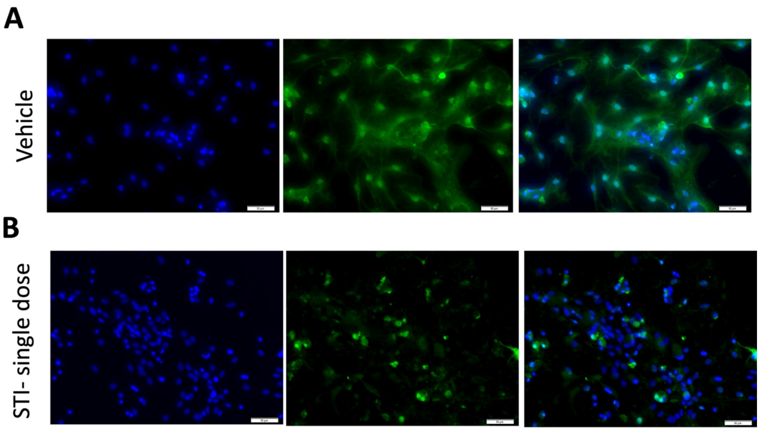


**Supplementary figure 2. Optimization of drug dose for STI.** (A) Vehicle. Experimental design: 2 x $10^5$ cells/mL of MGC were seeded in 24 well plates. After 48 hrs, the media was changed and replaced with FBS free media on day three and day six. The vehicle treatment with 300 µM concentration was given on day one and subsequently on day four. At the end on day 7 cells were stained with MAA lectin to see the expression of sialic acid. MAA recognizes α-(2,3)-linked sialic acid. and (B) After 48 hrs, cells were treated with STI at day 0 only and on day three cells were stained with MAA lectin to see the expression of α-(2,3)-linked sialic acid. Scale bar: 50 µm.

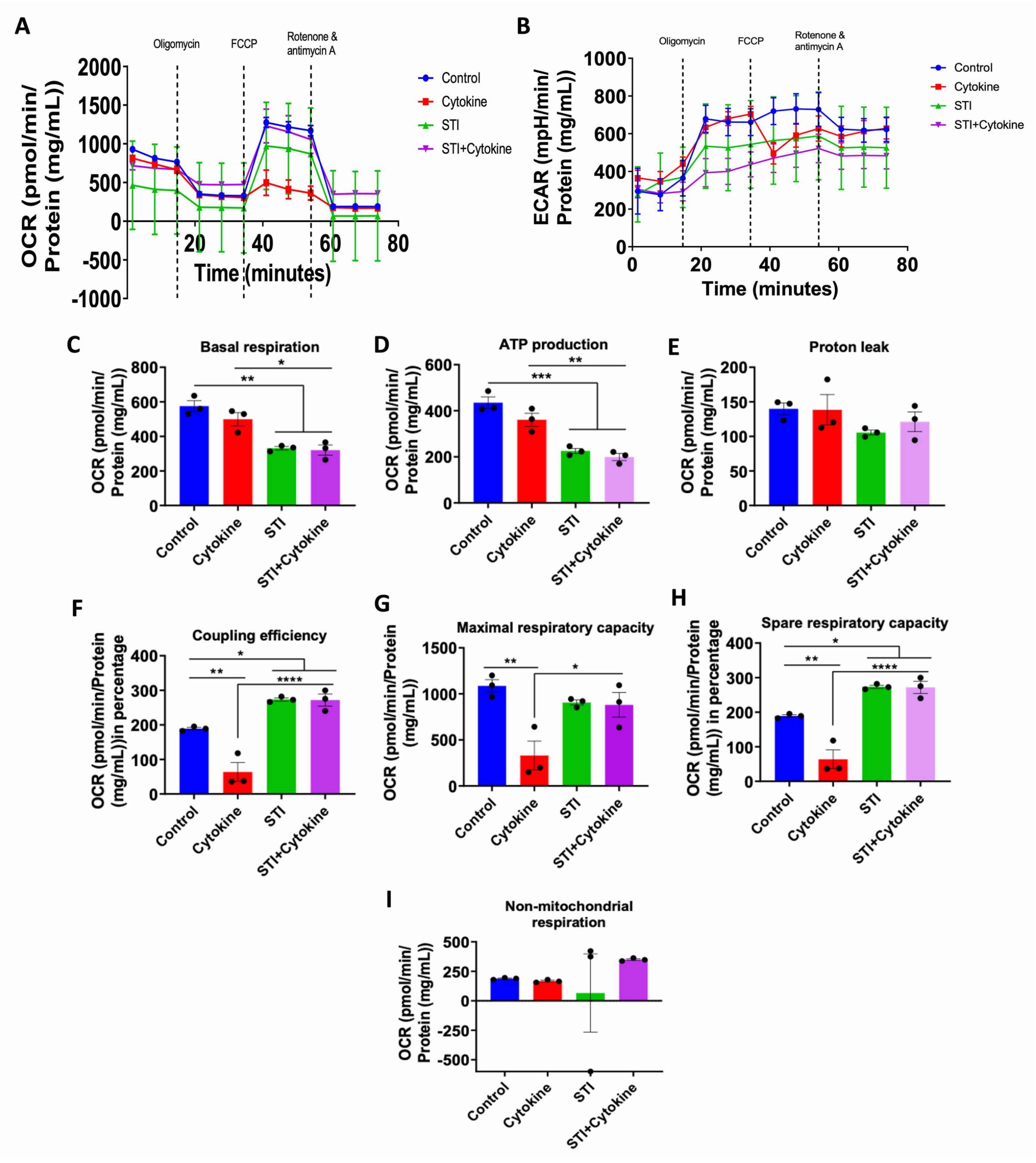


**Supplementary figure 3. A STI treatment reverts basal respiration, ATP production, proton leak, coupling efficiency, maximal respiratory capacity and spare respiratory capacity caused by cytokine treatment by day one.** After 48 hrs, MGC cells were treated with cytokine combination, with or without STI at day 0 only and on day three cells were undergone Cell Mito Stress assay. (A) OCR after the addition of three drugs (i.e. oligomycin, FCCP and Rotenone and antimycin A) sequentially. (B) Extracellular acidification rate (ECAR) after the addition of the above mentioned three drugs sequentially. (C-I) All parameters were calculated as a function of a cytokine combination and STI treatment. For this, total protein per well was calculated using a BCA protein quantification assay and data was normalised against it. Seven parameters namely, basal respiration, ATP production, proton leak, coupling efficiency, maximal respiratory capacity, spare respiratory capacity and non-mitochondrial respiration were measured and plotted as a bar graph. Data are represented as mean ± SEM, n=3. *$p$<0.05, **$p$<0.01, ***$p$<0.001, ****$p$<0.0001. One-way ANOVA followed by multiple comparison Tukey post hoc test was performed.

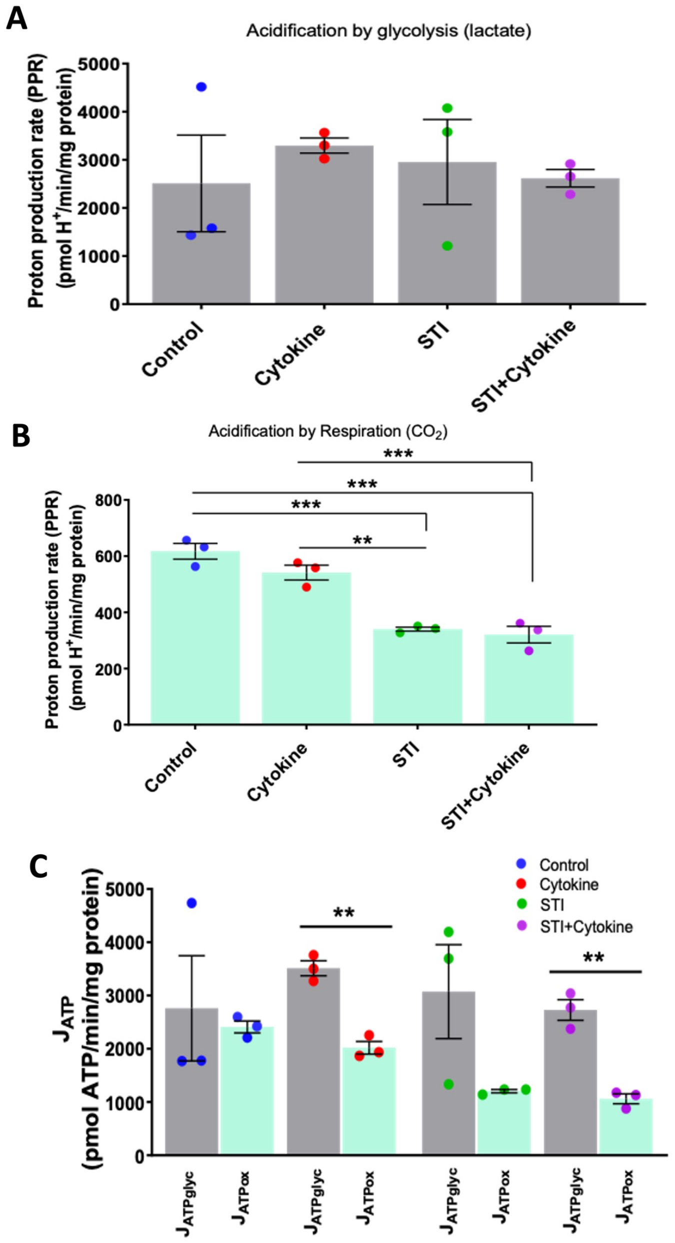


**Supplementary figure 4. Bioenergetics phenotype of mitochondrial function (at day one).** After 48 hrs, MGC cells were treated with cytokine combination, with or without STI at day 0 only and on day three cells were undergone Cell Mito Stress assay. (A) The rate of extracellular acidification caused by glycolysis due to lactate production. There was no difference in all the groups. (B) The rate of extracellular acidification due to respiration by $CO_2$ production. It was higher in control and cytokine treated group compared to the STI and STI + cytokine treated groups. However, there was no difference between the STI and the STI + cytokine treatment group in acidification by respiration. (C) Data from basal ECAR and OCR has been converted to the rate of ATP production by glycolysis and oxidation using formula. The rate of ATP production by glycolysis was not significant different between groups ($J_{ATPglyc}$). Under four treatments the rate of ATP production by glycolysis was higher than the rate of ATP production by oxidation. Data are represented as mean ± SEM, n=3. ***p*<0.01, ****p*<0.001, one-way ANOVA followed by multiple comparison Tukey *post hoc* test was performed.

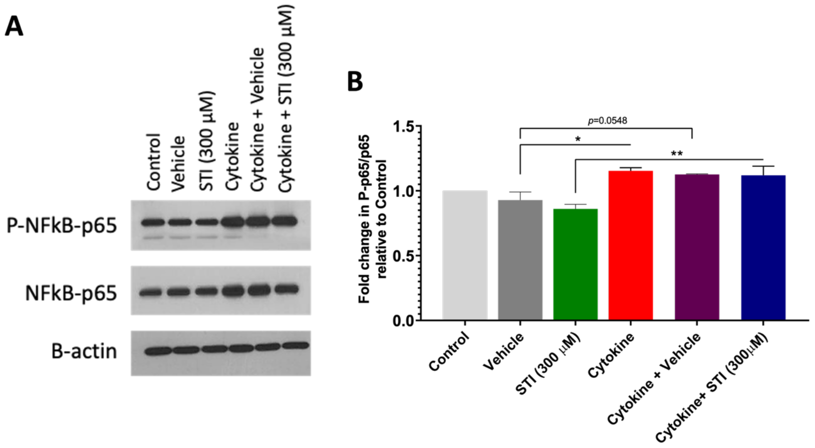


**Supplementary figure 5. Effect of STI on NFkB-p65 pathway in MGC under cytokine combination treatment.** After 48 hrs, cells were treated with STI with or without cytokine combination at day 0 only and on day three proteins were extracted and western blotting analysis was performed. (A) Western blots. (B) Quantification the intensity of the P-NFκB-p65 which was normalised to NFκB-p65 with respect to the control group. Alone STI treated group showed decreased expression of this pathway compared with all the groups with cytokine combination. Data are represented as mean ± SEM, n= three independent experiments. $^{**}p$<0.01, $^{*}p$<0.05; One-Way ANOVA, *post hoc* multiple comparison Tukey test.

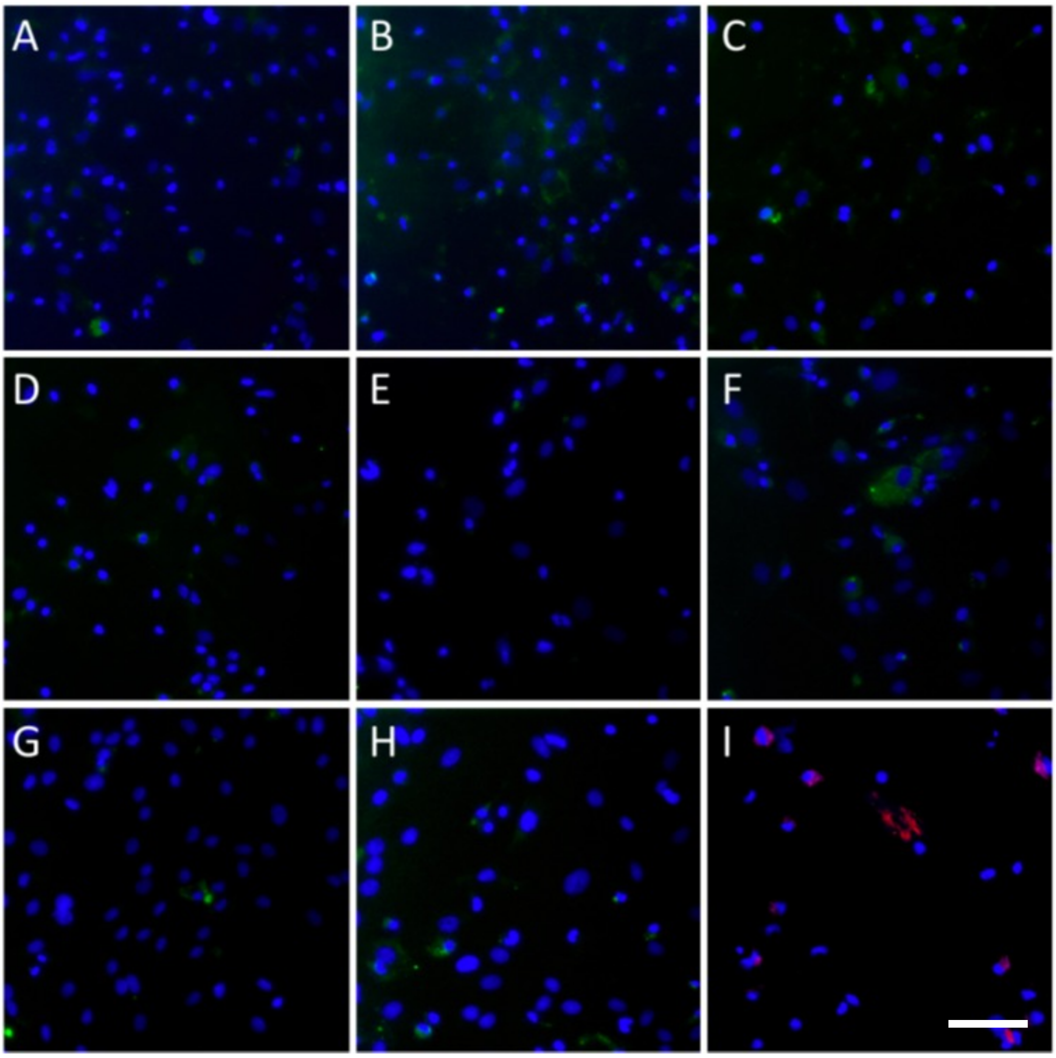


**Supplementary figure 6. Reduction in binding intensity after co-incubation of lectins with haptenic sugars demonstrates lectin binding is carbohydrate mediated.** All lectin staining was carried out in parallel in the presence of 100mM solution of the appropriate haptenic sugar. Competitive inhibition by these haptenic sugars resulted in a reduced binding intensity of the lectin to the glycans present in the MGC culture. The lectins and corresponding inhibitory carbohydrates were A) SNA-I in lactose, B) WFA in lactose, C) PNA in lactose, D) WGA in mannose, E) MAA in galactose, F) PHA-E in bovine IgG, and G) UEA-I in fucose. A control containing H) only TBS was conducted to determine the amount of auto-fluorescence. Finally, since the goal of this study was to do triple-staining for lectins, (GFAP and CD11b), I) secondary-antibody control was conducted to determine the amount of unspecific binding. Green = FITC, Blue = Hoechst, Red = Alexa Fluor 594, Purple = Alexa Fluor 647. Scale: 100 µm.

**Supplementary Table 1.** Lectin microarray Panel. Specificities of the lectins included in the array. Man: Mannose, GlcNAc: N-acetylgalactosamine, Gal: Galactose, LacNAc: N-acetyllactosamine, Sial: Sialic acid, Fuc: Fucose, a-Gal: α-Galactose.

| Lectin | Abbreviation | Specificity | Monosaccharide |
|---|---|---|---|
| Concanavalin A | ConA | a-Man, a-Glc | Man |
| Narcissus Pseudonarcissus | NPL | Terminal and internal Man | Man |
| Vicia faba lectin | VFA | a-Man, Glc, GlcNAc | Mannose |
| Allium sativum lectin | ASA | a-1,3-mannose | Mannose |
| Galanthus nivalis agglutinin | GNA | a-D-Man, terminal α-(1,3) mannose | Mannose |
| Triticum Vulgaris | WGA | (GlcNAc)n, sialic | GlcNac |
| Phytolacca americana | PWA | (GlcNAc)3 | GlcNAc |
| Griffonia simplicifolia II | BS-II | Terminal GlcNAc | GlcNAc |
| Lycopersicon esculentum | LEL | (GlcNAc)3 | GlcNAc |
| Solanum tuberosum | STL | (GlcNAc)3, LacNAc | GlcNac |
| Pseudomonas aeruginosa PA-I | PA-I | Gal | Gal |
| Datura stramonium | DSL | (β-1,4) linked N- acetylglucosamine oligomers, (GlcNAc)2-3, LacNAc | GlcNAc |
| Ricinus Communis Agglutinin, | RCA | a-Gal, Lac, LacNAc | Gal |
| Erythrina Cristagalli A | ECA | Gal, GalNac | GalNac |
| Phaseolus Vulgaris Agglutining | PHA E+L | oligos | Lac |
| Cicer arietinum | CAL | Fetuin, Lac, IgM | Lac |
| Maackia amurensis Lectin II | MAL-II | LacNAc | Gal |
| Sambucus nigra | SNA | a-2-6 sialic acid on LacNAc | Sialic, Lac |
| Maackia amurensis Lectin I | MAL-I | a-2,3-SialLacNac | Sial |
| Homarus americanus | HMA | LAg1 NeuNAc. LAg2: GalNAc | Sial, GalNAc |
| Ulex Europaea Aggl | UEA-I | a-1,2Fuc. L-fucose | Fuc |
| Pisum sativum | PSA | Fuca-1,6GlcNAc and a-Man | Fuc |
| Lotus tetragonolobus | LTL | Terminal a-Fuc, $Le^x$ | Fuc |
| Aspergillus orzyae | AOL | Fuc | Fuc |
| Aleuria aurantia lectin | AAL | α-1,6Fuc to N-acetylglucosamine, α-1,3Fuc to N-acetyllactosamine | Fuc |

| Lectin | Abbreviation | Specificity | Monosaccharide |
|---|---|---|---|
| Anguilla Anguilla agglutinin | AAA | D-Fucose, a-1,2Fuc, a-1,4Fuc | Fuc |
| Vicia villosa B4 | VVL | GalNAc | GalNac |
| Jacalin | JAC | T antigen | GalNac |
| Agaricus bisporus | ABL | b-Gal(1-3)GalNAc, T antigen | Gal |
| Amarantus Caudatus | ACA | T antigen, (Gala1-3GalNAca- Thr/Ser) | Gal |
| Maclura Pomifera | MPA | T antigen, aGalNAc | Gal |
| Psophocarpus tetragonolobus II | PT-II | a-1,2-fucosylated LacNAc | Gal |
| Griffonia simplicifolia I | BS-I | a-Gal | Gal |
| Euonymus Europaeus | EEA | Lac, blood groups B and H | Gal |
| Marasmium oreades agglutinin | MOA | Gal-α1,3Gal and Gal-α1,3Gal- β1,4GlcNAc | α-Gal |
| Helix pomatia | HPL | a-GalNAc terminal | GalNAc |
| Sophora japonica | SJA | GalNAc | GalNAc |
| Helix aspersa lectin | HAL | terminal N-acetyl-α-D- galactosamine | GalNAc |
| Soybean agglutinin | SBA | aGal-GalNAc | GalNAc |
| Peanut agglutinin | PNA | T antigen, Gal (b-1,3) GalNAc | Gal |
| Dolichos biflorus lectin | DBL | Terminal GalNAc | GalNAc |
| Wisteria floribunda | WFL | GalNAc | GalNAc |
| Salvia sclarea lectin | SSA | Terminal GalNac linked to serine | GalNAc |
| Psophocarpus tetragonolobus I | PT-I | a-GalNAc | GalNAc |
| Bauhinia Purpurea Lectin | BPL | GalNAc | GalNAc |
| Lectin from Arachis hypogaea (peanut) | AHP | T antigen, Gal(β-1,3) GalNAc | Gal |
| Chlostridium fragile lectin | CFL | N-acetyl-D-galatosamine | GalNAc |
| Phaseolus lunatus (Lima bean) agglutinin | LBA | GalNAc, GalNAcα-(1,3) [L-fuc α(1,2)]Gal | GalNAc |

### **Supplementary Table 2.** Lectin immunostaining panel.

| Abbreviation | Lectin | Concentration Used | Origin | Binding Specificity | Inhibitory Carbohydrate (Haptenic sugar) (100mM) |
|---|---|---|---|---|---|
| A.SNA-I | Sambucus nigra Isolectin-I | 20 µg/mL | Sambucus nigra (Elderberry) | Neu-α-(2 → 6)-Gal(NAc)-R | Lactose |
| B. WFA | Wisteria Floribunda agglutinin | 30 µg/mL | Wisteria Floribunda (Japanese wisteria) | terminal α- or β-linked GalNAc, lactose, Gal, chondroitin sulfate | Lactose |
| C. PNA | Peanut agglutinin | 20 µg/mL | Peanut (Arachis hypogaea) | Gal-β-1→ 3Gal(Nac)-R | Lactose |
| D. WGA | Wheat germ agglutinin | 20 µg/mL | Wheat germ | Glc(Nac)-β-(1→ 4)-Glc(Nac)-(1→4)>-β-Glc(Nac) | Mannose |
| E. MAA | Maackia amurensis agglutinin | 20 µg/mL | Maackia amurensis (Amur maackia) | Neu-α-(2 → 3)-Gal-β-(1 → 4)-GlcNAc-R, $SO_4^{2-}$-3-Gal-β-(1 → 4)-GlcNAc-R | Galactose |
| F. PHA-E | PHASEOLUS VULGARIS LECTIN | 10 µg/mL | Red kidney bean | Gal-β-(1→4)-GlcNAc-β-(1→2)-Manα | Bovine IgG |
| G. UEA-I | Ulex europaeus agglutinin | 20 µg/mL | Gorse, Furze | L-Fu-α-(1→2)-Gal-β-(1→4)-Glc(Nac)-β(1→6)-R | Fucose |
| H. GNA | Galanthus nivalis agglutinin | 20 µg/mL | Snowdrop bulbs | α-1,3 mannose residues | Mannose |
| I. DSL | Datura stramonium lectin | 20 µg/mL | Datura stramonium (Jimson weed) | (β-1,4) linked N-acetylglucosamine | GlcNAc |
| J. RCA-I | Ricinus communis I | 10 µg/mL | Castor bean | galactose or N-acetylgalactosamine residues | Galactose |

### **Supplementary Table 3. Summary of the glycan expressions analyzed by lectin staining at day one.**

| Abbreviation | Lectin | Carbohydrate specificity | Output |
|---|---|---|---|
| A.SNA-I | Sambucus nigra isolectin-I | Neu-α-(2 → 6)-Gal(NAc)-R | The α-2,6-sialic acid moieties were associated with microglia than with astrocytes, and quantification showed a higher signal in the cytokine-combination treated group (1.29-fold). LPS did not show any change compared to other groups. |
| B. WFA | Wisteria Floribunda agglutinin | Terminal α- or β-linked GalNAc, lactose, Gal, chondroitin sulfate | The α or β-GalNAc residues were associated with astrocytes in the control and cytokine treated groups. However, LPS treatment showed lower WFA binding ($p<0.01$). |
| C. PNA | Peanut agglutinin | Gal-β-1→ 3Gal(Nac)-R | The β-(1-3)-GalNAc residue, showed increased binding with astrocytes and microglia upon cytokine combination (1.24-fold), whereas this decreased after LPS treatment ($p<0.0549$). |
| D. WGA | Wheat germ agglutinin | Glc(Nac)-β-(1→ 4)-Glc(Nac)-(1→4)>-β-Glc(Nac) | The β-(1-4)-GlcNAc residue, showed increased binding with astrocytes and microglia upon cytokine combination (0.91-fold), whereas this decreased after LPS treatment (1.95-fold). |
| E. MAA | Maackia amurensis agglutinin | Neu-α-(2 → 3)-Gal-β-(1 → 4)-GlcNAc-R, $SO_4^{2-}$-3-Gal-β-(1 → 4)-GlcNAc-R | The α-2,3-sialyation was strongly expressed on microglia, and upon LPS treatment the expression was reduced ($p<0.01$). |
| F. PHA-E | Phaseolus vulgaris lectin | Gal-β-(1→4)-GlcNAc-β-(1→2)-Manα | No difference between control and cytokine-combination treated samples; however, LPS treatment reduced ($p<0.05$) expression level of GlcNAc residues. |
| G. UEA-I | Ulex europaeus agglutinin | L-Fu-α-(1→2)-Gal-β-(1→4)-Glc(Nac)-β(1→6)-R | UEA-I lectin showed binding for fucosylation on both astrocytes and microglia and there was no difference between control and, cytokine-combination and LPS treated groups. |
| H. GNA | Galanthus nivalis agglutinin | α-1,3 mannose residues | The expression of α-1,3 mannose on both astrocytes and microglia, and the LPS treated group showed higher expression at day one (7.34-fold). |
| I. DSL | Datura stramonium lectin | (β-1,4) linked N-acetylglucosamine | The β-(1-3)-GalNAc residues were associated with microglia and astrocytes in MGC, and there was no significant difference between control, cytokine combination and LPS treated groups. |
| J. RCA-I | Ricinus communis I | Galactose or N-acetylgalactosamine residues | GalNAc residues expressed on astrocytes and microglia and was higher (1.92-fold) in the cytokine-combination-treated group. |
